# Malaria vector diversity, transmission, and insecticide resistance, in island communities along the Volta Lake in southern Ghana

**DOI:** 10.1101/2025.01.23.634470

**Authors:** Abena Ahema Ebuako, Christopher Mfum Owusu-Asenso, Abdul Rahim Mohammed, Isaac Kwame Sraku, Yaw Akuamoah-Boateng, Cornelia Appiah-Kwarteng, Akua Obeng Forson, Patrick Ferdinand Ayeh-Kumi, Yaw A. Afrane

## Abstract

Island communities along the Volta Lake in southern Ghana present unique challenges for malaria control, characterized by high transmission rates, limited vector control measures, and isolated ecosystems. This study assessed the malaria vector diversity, seasonal abundance, transmission, and insecticide resistance status of malaria vectors in these communities to inform effective control strategies. Mosquitoes were collected from three island communities (Tuanikope, Allorkpem, and Pediatorkope) using human landing catches, light traps, and prokopack aspirators during the dry and rainy seasons. Morphological and molecular techniques were used to identify mosquito species, determine blood meal sources, and detect insecticide resistance mutations. Sporozoite infections and entomological inoculation rates (EIRs) were also quantified. A total of 25,079 mosquitoes from four genera were collected; Culicine = 88.19%, Anopheline = 8.89%. Mansonia = 2.24%. The Anophelines predominantly comprised *Anopheles gambiae* s.l. 1,911/2,230 (85.70%), followed by *An*. *pharoensis* 236/2,230 (10.58%) and *An. rufipes* 83/2,230 (3.72%). Indoor biting and resting densities were high across sites and seasons, with sporozoite-positive mosquitoes more frequently found indoors. Blood meal analysis revealed a strong anthropophilic feeding pattern (HBI = 80%). Annual EIRs ranged from 37.40 (ib/m/y) to 100.08 (ib/m/y). Low frequencies of insecticide resistance mutations (Vgsc-1014F, Vgsc-1014S, ACE-1, and Vgsc-1575Y) were observed. The study indicates high indoor biting and resting densities of *Anopheles* mosquitoes. High sporozoite rate along with low resistance mutation frequencies observed, emphasize the urgent need for continuous resistance monitoring and the implementation of targeted vector control strategies in these hard-to-reach island communities.

## Introduction

In Ghana, the *Anopheles gambiae* sensu lato and *Anopheles funestus* are the predominant malaria vector species [1–3]. The distribution of these vectors varies with changes in climate and ecological conditions coupled with other factors such as land usage patterns and vector-host interactions [2,4]. Island communities present a distinct setting for malaria transmission characterized by isolated ecosystems and limited access to healthcare resources. Despite their isolation, Island populations remain vulnerable to malaria due to environmental suitability for mosquito breeding and limited implementation of preventive measures [5,6]

Previous studies in Africa have shown that malaria epidemiology within island ecosystems is characterized by a diverse array of vector species, each exhibiting unique biology, behaviour, and vectorial capacity [7–10]. Malaria vectors in Island communities may undergo specific adaptations to the local ecosystem [9]. These adaptations include changes in behaviour, physiology, or genetics, reflecting the vectors’ ability to thrive in Island conditions [7]. Furthermore, seasonal variations in temperature, rainfall, and humidity may affect the biology of malaria vectors [11]. Islands experience unique climate patterns that may also influence vector abundance, longevity, and overall transmission intensity.

Whilst several studies have reported on the malaria vector bionomics and insecticide resistance status across Ghana’s various ecological zones [3,12–14], there is a paucity of data on the bionomics and insecticide resistance status of the malaria vectors in Island communities in Ghana. These Island communities face challenges distinct from those on the mainland, including a lack of a structured vector control program [15,16], limited access to essential social amenities, and geographical barriers that hinder effective vector surveillance, malaria transmission studies, and preventive research in these isolated communities [17]. Island communities are also affected by several other factors, such as environmental conditions, human behavior, and climate change, all of which have an impact on malaria transmission [8,18]. These limitations may translate to a high rate of malaria transmission as a result of the increased density of the disease vectors.

To efficiently control malaria vectors and ultimately eliminate malaria in Ghana by 2030, it is important to have up-to-date information on local vector bionomics and insecticide resistance to inform the National Malaria Elimination Program to make timely changes to vector control strategies. This study assessed the malaria vector diversity, seasonal abundance, transmission, and insecticide resistance status of malaria vectors in Island communities on the Volta Lake in Southern Ghana.

## Materials and methods

### Study sites

The study was conducted in three Island communities; Tuanikope, Pediatorkope, and Allorkpem on the Volta Lake, in the Dangme East District of Greater Accra in Southern Ghana (Fig 1). The inhabitants of these communities are usually fish farmers. The presence of abandoned canoes and backyard farming activities collects pockets of potential mosquito breeding habitat which could support the breeding of mosquitoes. These are hard-to-reach Island communities that lack structured malaria control interventions such as IRS and the use of LLINs. In an earlier study, it was realized that these island communities have not received vector control intervention for over eight years [15,16]Malaria transmission occurs all year round, with seasonal peaks during the rainy season [15]. The communities lack electricity, causing the inhabitants to spend longer hours outdoors, which could facilitate man-vector contact. These two island communities lie in the coastal savannah ecological zone in Ghana which has a bimodal rainfall pattern. The main rainy season is from April to June and the main dry season is from January to March.

**Fig 1:**
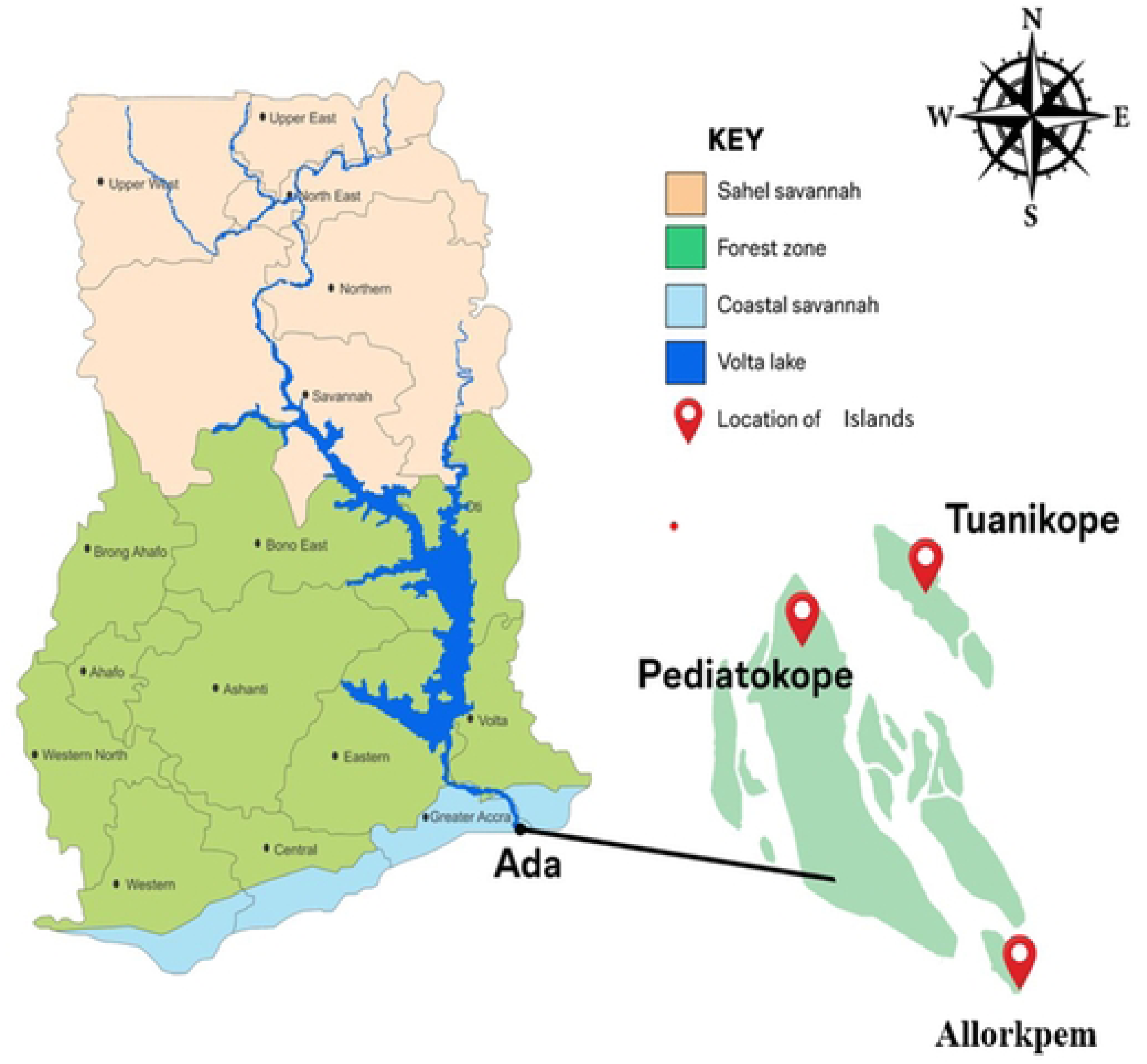
Map of Ghana showing study sites.

### Adult *Anopheles* mosquito collections

At each sampling site, sixteen houses were randomly selected, and sampled during the dry (25^th^ April – 6^th^ May, 2023) and rainy seasons (27^th^ July – 9^th^ August, 2023). Sampling was done simultaneously indoors and outdoors in each house. In the dry season, *Anopheles* mosquitoes were collected for four consecutive nights, and three nights in the rainy season (Four houses per day). Indoor and outdoor host-seeking mosquitoes were collected using the Human Landing Catch (HLC) technique [19] and light traps (John W. Hock Company, Gainesville, Florida, USA), whereas resting mosquitoes were collected using the prokopack aspirators (John W. Hock, Gainesville, FL). Each study site was divided into 4 sections to ensure a fair representation of the mosquito population at the study site.

Individuals aged 18 years and above who voluntarily opted to collect mosquitoes using the HLC method were trained to expose their lower limbs as bait to attract mosquitoes. Briefly, volunteers sat in the dark with their lower limbs exposed and with the aid of a flashlight, located and collected the host-seeking mosquitoes with a collection tube when they landed in search of a blood meal. Prior to training and mosquito sampling, all adult volunteers participating in the HLC study provided written informed consent. Volunteers received a copy of their signed consent form, while another copy was securely stored in a locked cabinet within the Department of Medical Microbiology at the University of Ghana Medical School. Individuals who consented to partake in this study were given prophylaxis before the HLC collections. Independent staff supervised and regularly walked between different groups throughout the night, for quality control of collectors placed inside and outside selected houses[20].

The collection of mosquitoes using light traps took place in houses where the Human Landing Catch (HLC) method was not being used. Three houses were randomly selected for four nights and light traps were placed at a height of one meter from the ground of each house and baited with a mixture of sugar, yeast, and water to produce carbon dioxide to attract mosquitoes [21]. They were set up both inside and outside households at 18:00h and retrieved at 06:00h the following morning.

The prokopack aspirator was used to hoover the ceiling, under tables, walls and all possible resting surfaces of mosquitoes to collect any resting mosquitoes. Five houses were randomly sampled for four nights between 5:30 h to 7:00 h during both seasons. Households were advised the night before sampling to refrain from opening their doors and windows upon waking the next morning to prevent resting mosquitoes from exiting their rooms.

### Morphological identification of *Anopheles* mosquitoes

All adult mosquitoes collected were knocked out by exposure to chloroform. Samples were sorted into the different genera (*Anopheles*, *Mansonia*, *Aedes,* or *Culex*) and sex based on the identification keys by Gillies & Coetzee, (1987). Female *Anopheles* mosquitoes were additionally categorized based on their gonotrophic status as unfed, blood-fed, half-gravid, or gravid. All samples were individually packed on silica gel in Eppendorf tubes and labeled with the study site and date of collection [22]. Resting blood-fed *Anopheles* mosquitoes were placed separately in 1.5 ml Eppendorf tubes with 100% ethanol and then labeled. All samples were transported to the Parasitology laboratory of the Department of Medical Microbiology, University of Ghana.

### Molecular identification of vector mosquitoes

A sub-sample from the total *An. gambiae* s.l. collected over the study period was selected according to study site, season, and location (indoor or outdoor) in a proportion of ten percent (10%) and used to discriminate the species. Individual mosquito legs were cut and used as DNA templates to undertake PCR for discriminating the sibling species using the method of Scott *et al* [23].

### Detection of sporozoites

Genomic DNA extracted from the head and thoraces of sub-selected mosquitoes used for speciation were used to detect the presence of *Plasmodium* sporozoites by PCR as described by Echeverry *et al*., [24].

### Blood meal analysis in mosquitoes

To identify the origin of the blood meals, genomic DNA was extracted from the abdomen of blood-fed *Anopheles* mosquitoes using the ZR DNA MicroPrep kit, followed by PCR amplification of the mitochondrial cytochrome b gene with one universal reverse and five animal-specific forward primers (human, cow, goat, pig, dog) [25]. Positive controls of selected vertebrate s and a negative control were included in the PCR reaction. The PCR products were electrophoresed on a 2 % agarose gel. The results were interpreted using the following band sizes: 453bp (Pig), 334bp (Human) 132bp (Goat), 680bp (Dog), 561bp (Cow).

### Determination of insecticide-resistant mutations in *An*. *gambiae* s.l

Genomic DNA extracted from the heads and thoraces of the indoor and outdoor mosquito samples was used to detect the presence of insecticide resistance mutations using TaqMan SNP genotyping probe-based assays [26]. These markers included Vgsc-1014F, Vgsc-1014S, Ace1-119S and Vgsc-1575Y mutations.

### Data management and analysis

Descriptive analysis was done to visualize resistant allele frequencies, and mosquito species composition from the selected sites using graphs and tables. Chi-square analysis was used to determine if the data on two categories were associated. The number of female mosquitoes collected using the Prokopack aspirator was used to calculate the resting density of Anopheline mosquitoes per site. The ratio of blood-fed mosquito samples that had drawn blood from people to the total number of mosquitoes screened for blood meal sources yielded the Human Blood Index (HBI). By dividing the total number of mosquitoes tested for infection by the number of sporozoite-positive mosquitoes, the sporozoite infection rate (SIR) was calculated. The total number of mosquitoes collected by HLC divided by the total number of collectors per total number of nights collected was used to compute the Human Biting Rate (HBR). The product of the sporozoite rate and the human biting rate was used to compute the Entomological Inoculation Rate (EIR). Allele frequencies of resistance gene markers in the vector populations at each site were calculated using Hardy-Weinberg equilibrium (HWE). All statistical analyses were done in and STATA/IC 14.1.

## Results

### Abundance and seasonal distribution of malaria vectors

Overall, 25,079 mosquitoes were collected belonging to four different genera during the sampling period. Mosquitoes collected in the dry season (n= 17,638; 70.33%) were over twice as many as mosquitoes collected in the rainy season (n = 7,441; 29.67%). Out of the total mosquitoes sampled, 22,116 (88.19%) belonged to the Culicines, 2,230 (8.89%) Anophelines, and 563 (2.24%) were Mansonia. The majority of Anophelines caught belonged to the *Anopheles gambiae* s.l. 1,911/2,230 (85.70%) followed by *An*. *pharoensis* 236/2,230 (10.58%) and *An. rufipes* 83/2,230 (3.72%) (Table 1).

**Table 1:** Abundance and spatiotemporal distribution of mosquito genera and *Anopheles* species.

| Mosquitoes | Dry Season |  |  |  | Rainy Season |  |  |  |
| --- | --- | --- | --- | --- | --- | --- | --- | --- |
|  | Allorkpem | Pediatorkope | Tuanikope | Total | Allorkpem | Pediatorkope | Tuanikope | Total |
| *An. gambiae* s.l. | 706 | 252 | 592 | 1,550 | 155 | 76 | 130 | 361 |
| *An. pharoensis* | 2 | 1 | 46 | 49 | 8 | 8 | 18 | 34 |
| *An. rufipes* | 80 | 84 | 30 | 194 | 28 | 7 | 7 | 42 |
| Culicine | 5,792 | 5,350 | 4,605 | 15,747 | 2,691 | 1,173 | 2,505 | 6,369 |
| Mansonia | 84 | 3 | 11 | 98 | 34 | 166 | 435 | 635 |
| **Total** | 6,691 | 5,746 | 5,296 | **17,733** | 2,924 | 1,434 | 3,097 | **7455** |

Anopheline diversity observed in this study was higher in the dry season [ Dry = 80.40% (*An. gambiae* s.l. n = 1550/1793 (86.45%), *An*. *rufipes* n = 194/1793 (10.82%), *An*. *pharoensis* n = 49/1793 (2.73%)]; Rainy = (*An. gambiae* s.l. n = 361/437 (82.61%), *An*. *rufipes* n = 42/437 (9.61%), *An*. *pharoensis* n = 34/437 (7.78%)], (Table 1). Among the studied locations, An. gambiae s.l. was the most abundant in Allorkpem. [dry (706/1550); rainy (155/361)], followed by Tuanikope [dry (592/1550); rainy (130/361)] and Pediatorkope [dry (252/1550); rainy (76/36)] in both seasons (Table 1).

Overall HLC [1,814 (94.92%)], was the most effective trapping method for sampling *An. gambiae* s.l., followed by light trap [73 (3.82%)], and prokopack [24 (1.26%)].

### Indoor and Outdoor Biting Activity of *Anopheles* mosquitoes

*Anopheles gambiae* s.l. collected from each study site demonstrated strong endophagic behavior (Allorkpem, 65.51% n = 564/861; Pediatorkope, 70.12% n = 230/328 and Tuanikope, 50.14% n = 362/722). However, *An. rufipes* (Allorkpem, 55.15%; Pediatorkope, 64.29% and Tuanikope, 56.78%) and *An. pharoensis* (Allorkpem, 55.10%; Pediatorkope, 52.94% and Tuanikope, 54.22%) preferred feeding outdoors (exophagy) (Table 2). The indoor human biting rate (HBR) of *An. gambiae* s.l. was higher than outdoor in Allorkpem [dry (13.78 vs. 7.41 mosquitoes/person/night) (m/p/n); rainy (3.88 vs. 2.33 m/p/n)] and Pediatorkope [dry (4.97 vs. 2.38 m/p/n); rainy (2.21 vs. 0.78 m/p/n)] for both seasons. In contrast to the other sites, HBR in Tuanikope was higher outdoors than indoors during the dry season (9.16 vs. 8.34 m/p/n). Whilst in the rainy season, indoor HBR of *An. gambiae* s.l. remained higher than outdoors (2.63 vs. 2.42 m/p/n).

**Table 2:**
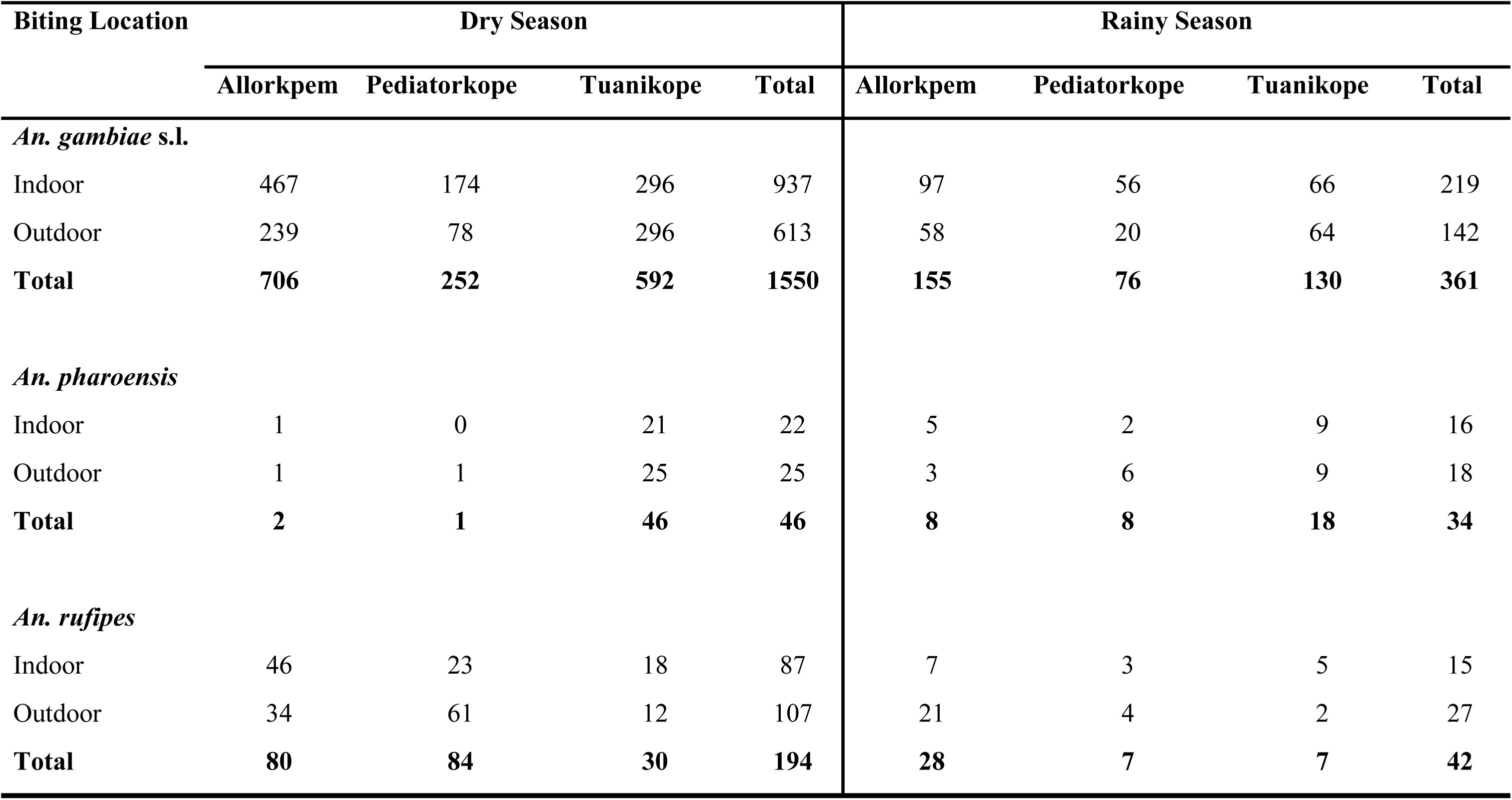
Biting location of *Anopheles* mosquitoes.

### Biting patterns of *Anopheles gambiae* s.l

*Anopheles gambiae* s.l. had a preference for late evening/classical biting (LE: 22:00 – 4:00 h) (66.2%; 1134/1814) followed by early morning (EM: 4:00 – 6:00 h) (21.11%; 383/1814), and early evening (EE: 18:00 – 22:00 h) (16.37%; 297/1814). The pattern remained the same for indoors [LE: (67.47%, 726/1076); EM: (19.52%, 210/1076); EE: (13.01%, 140/1076)] and outdoors biting [LE: (55.28%, 408/738); EM: (23.44%, 173/738), EE: (21.27%, 157/738)].

The man-biting activity of *An. gambiae* s.l. started from dusk to dawn both indoors and outdoors in all study sites. In Allorkpem, the biting activity was bimodal in both seasons with the highest biting activity observed after midnight (01:00h - 02:00 h) [dry = 2.1 mean bites/person/hour(m/p/h); rainy = 0.8 m/ p/h], (Fig 2A). The biting activity of *An. gambiae* s.l. decreased steadily in the early morning, till 05:00-06:00 h (0.8 m/ p/h) in the dry season, (Fig 2A). However, there was a sharp increase in early morning bites with a peak biting activity at 05:00-06:00 h (1.8 m/ p/h) in the rainy season, (Fig 2A).

**Fig 2:**
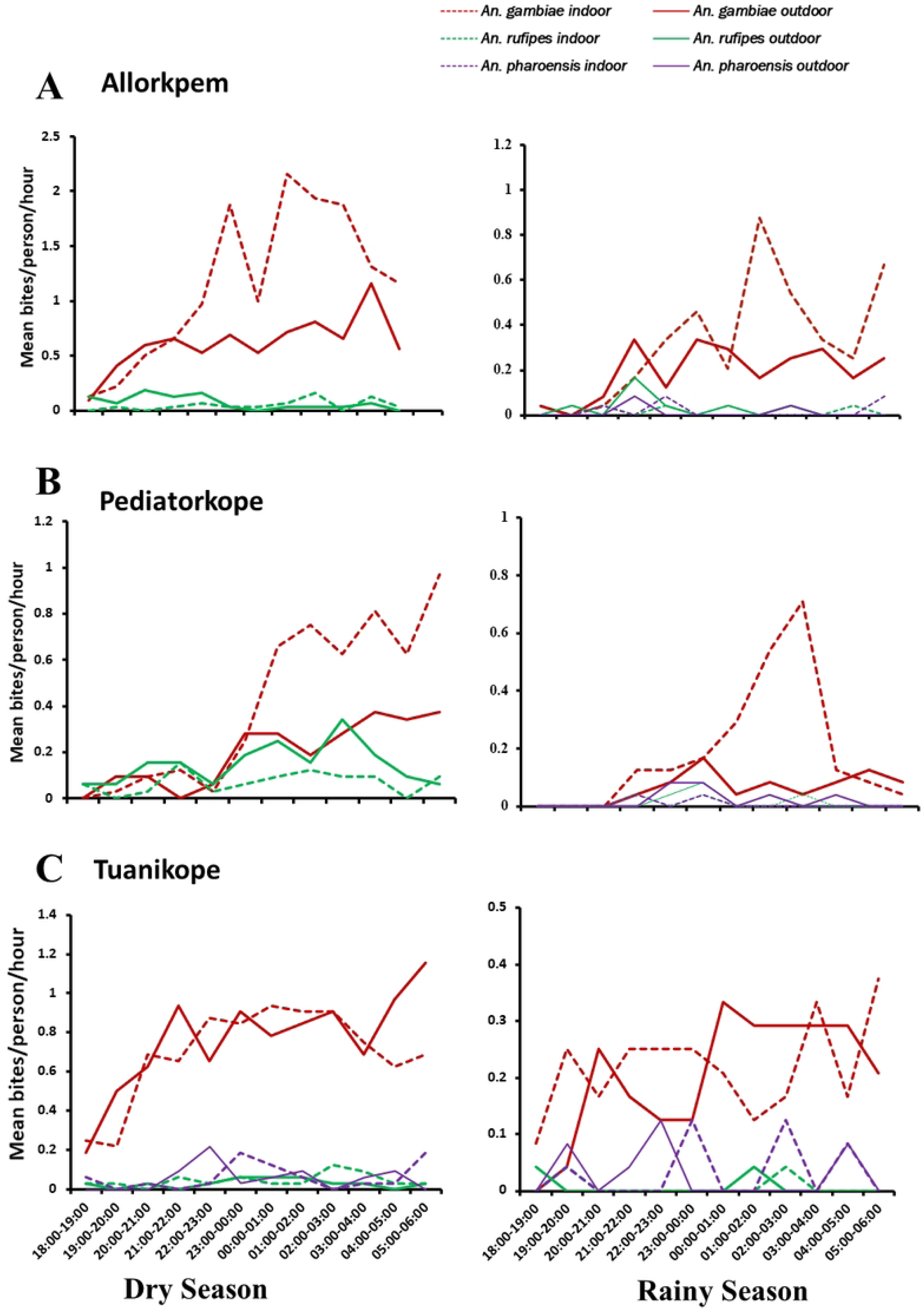
Hourly mean biting rate in (A):Allorkpem, (B):Tuanikope and (C):Pediatorkope.

In Pediatorkope, *An*. *gambiae* s.l. showed multiple peaks after midnight with the highest biting activity at 05:00 - 06:00 h indoors during the dry season (0.9 m/ p/h), (Fig 2B). In the rainy season, biting activity peaked at 02:00 - 03:00 h (0.7 m/ p/h) indoors. This surge in biting activity was followed by a sharp decline, (Fig 2B).

In Tuanikope the biting activity of *An. gambie* s.l. peaked at different hours of the night in both seasons and locations. A major outdoor biting activity was observed in the early morning (05:00-06:00 h) during the dry season (1.16 m/ p/h), (Fig 2C). However, in the rainy season high indoor biting activity was observed (0.36 m/ p/h) at the same time (Fig 2C).

### Resting densities of *Anopheles gambiae* s.l

Out of the 1,911 *An*. *gambiae* s.l collected; 24 resting mosquitoes were caught using the prokopack aspirator. Overall, majority of the resting mosquitoes were collected during the rainy season accounting for 52.17 % (13/24), compared to 47.83 % (11/24) collected during the dry season. Majority of the mosquitoes were sampled indoors (52.17 % (13/24)) whiles 47.83 % (11/24) were collected outdoors. During the dry season, the highest indoor resting density was recorded in Tuanikope (0.25, n = 5/11). However, the highest resting density during the rainy season was recorded outdoors in Tuanikope (0.4, n =6/13) (Fig 3).

**Fig 3:**
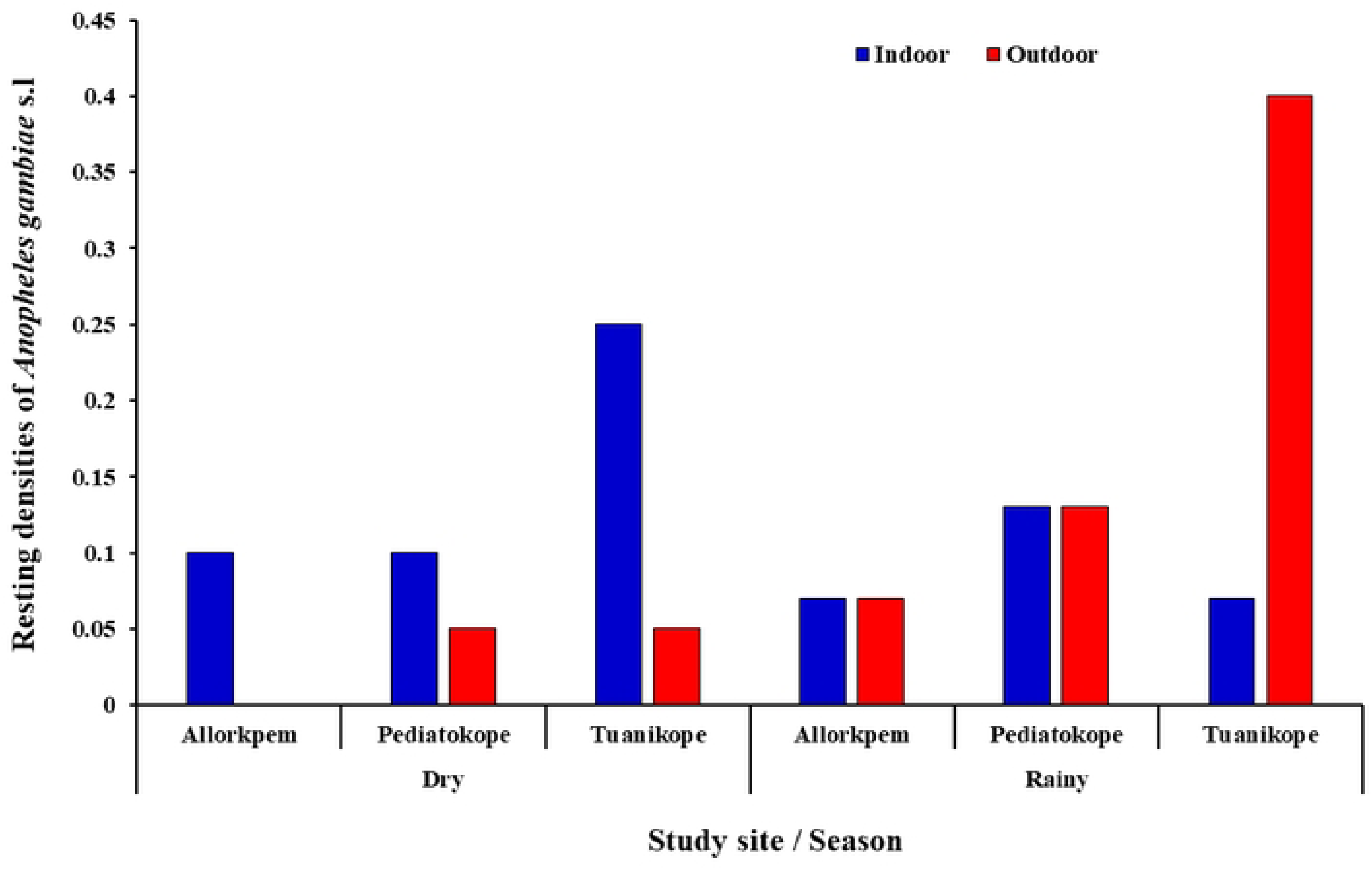
Resting densities of *An. gambiae* s.l. collected from Island communities.

### Species discrimination of the *An. gambiae* complex

From the total *An*. *gambiae* s.l. collected across all sites, a subsample of 862 were randomly selected and used to distinguish between the sibling species. This resulted in 62.88% (n = 538/862) *An. gambiae* s.s., *An. coluzzii* 37.04% (n = 316/862), and hybrid 0.94% (n = 8/862), (Fig 4). The species composition and spatial distribution exhibited significant differences across seasons (χ^2^ = 124.3045.92, df = 2, *P* = 0.000), location of collection (χ^2^ = 39.8512, df = 2, *P* = 0.000), and study sites (χ^2^ = 149.4988, df = 4, *P* = 0.000). Overall, *Anopheles gambiae* s.s. dominated mosquito populations across island communities, except in Pediatorkope, where *An. coluzzii* were more abundant: Allorkpem = [ (*An*. *gambiae* s.s. = 257/323; *An*. *coluzzii* = 66/323); Pediatorkope = (*An*. *gambiae* s.s. = 36/158; *An*. *coluzzii* = 122/158); Tuanikope = (*An*. *gambiae* s.s. = 244/373; *An*. *coluzzii* = 129/373)] (χ^2^ = 283.27, df = 6, *P* = 0.000)

**Fig 4:**
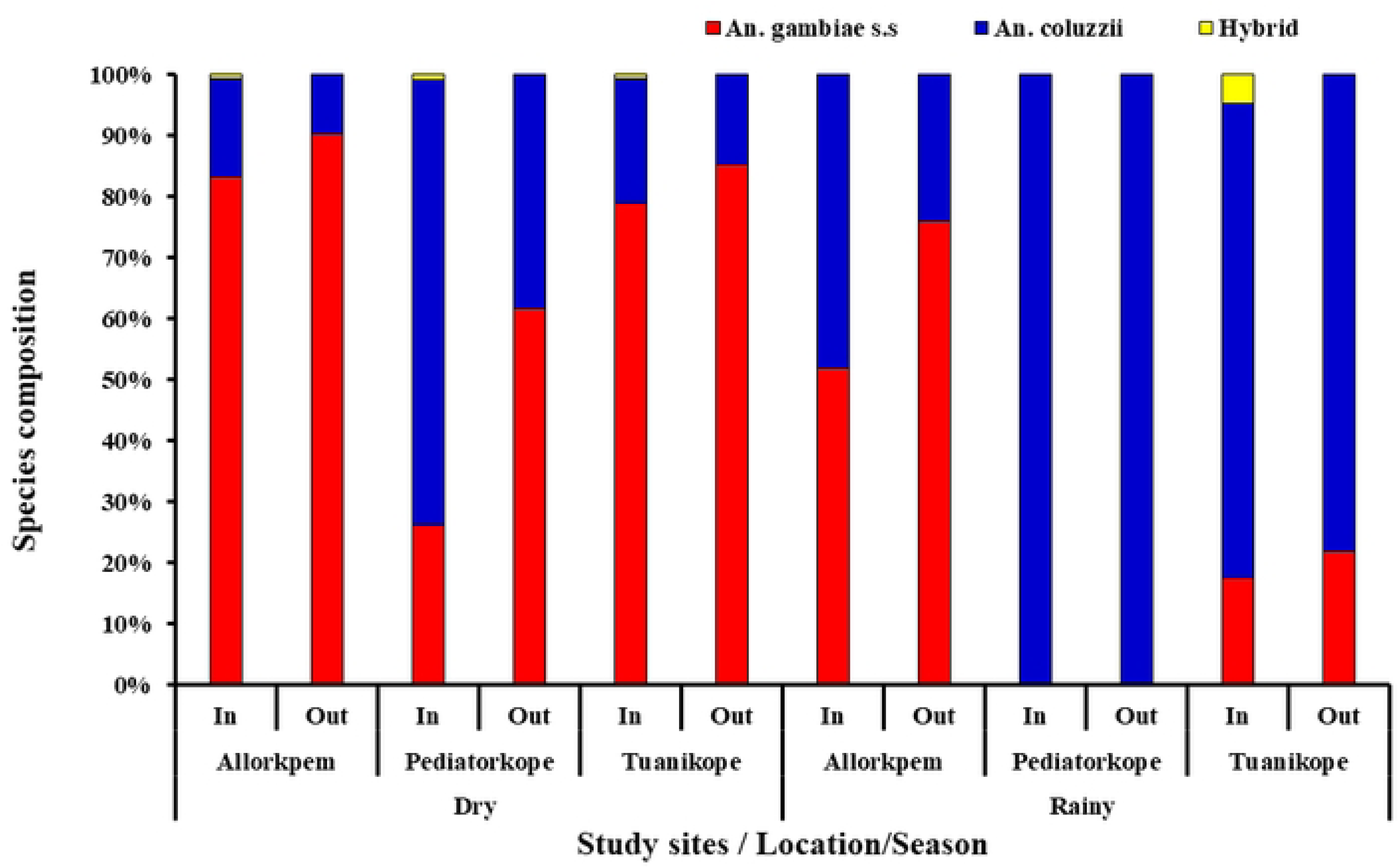
Species discrimination in the *Anopheles gambiae* s.l.

### Sporozoite infectivity rates in the sampled vectors

In addition, all 862 sub-samples that were genotyped to species were used to determine the presence of sporozoites. Thirty-one (31) samples [dry = 15/640; rainy = 16/222] (χ^2^ = 2.2485, df =1, *P* = 0.134) were positive for *P. falciparum* CSP. Infected mosquitoes were identified from all three island communities: Allorkpem: (0.02), n = 8/326, Pediatorkope: (0.04), n = 6/160 and Tuanikope: (0.05), n = 17/376, with no significant variation in sporozoite rate (χ^2^ = 0.1690, df = 1, *P* = 0.681). Though more CSP-positive *An*. *gambiae* s.l mosquitoes were collected indoors (0.61), n = 19/31 than outdoors (0.39), n = 12/31, the difference was not significant (χ^2^ = 0.0937, df = 1, *P* = 0.78). Overall, majority of the infected mosquitoes were *Anopheles coluzzii* (n = 20/31) and were collected during the classical biting time (12 am - 3 am).

### Entomological inoculation rates in the sampled vectors

Overall, Tuanikope had the highest estimated annual EIR of 100.08 (ib/m/y) followed by Allorkpem [65.94 (ib/m/y)] and Pediatorkope [37.40 (ib/m/y)]. In the dry season, an unprotected individual living in Allorkpem was most exposed to an infective bite from *Anopheles* mosquito with an EIR of 0.26 ib/m/n. Tuanikope 0.19 (ib/m/n) followed as the next highest risk area, with Pediatorkope 0.11 (ib/m/n) having the lowest exposure among the three locations. Conversely, in the rainy season, individuals in Tuanikope were more exposed to infective bites with an EIR of 0.28 ib/m/n followed by Pediatorkope 0.11ib/m/n and Allorkpem 0.07 ib/m/n. (Table 3)

**Table 3:** Entomological Inoculation rate for Island communities.

| Study sites | Season | HBR(b/p) | SPR | EIR(ib/m/n) | EstEIR/y(ib/m/y) |
| --- | --- | --- | --- | --- | --- |
| Allorkpem | Dry | 10.59 | 0.03 | 0.26 | 96.63 |
|  | Rainy | 3.1 | 0.02 | 0.07 | 27.16 |
|  | **TOTAL** | **7.38** | **0.02** | **0.18** | **65.94** |
| Pediatorkope | Dry | 3.67 | 0.03 | 0.09 | 33.49 |
|  | Rainy | 1.48 | 0.08 | 0.11 | 40.52 |
|  | **TOTAL** | **2.73** | **0.04** | **0.10** | **37.40** |
| Tuanikorpe | Dry | 8.75 | 0.02 | 0.19 | 70.26 |
|  | Rainy | 2.52 | 0.11 | 0.28 | 101.18 |
|  | **TOTAL** | **6.08** | **0.05** | **0.27** | **100.08** |

HBR=man biting rate, SPR=sporozoite rate, EIR= entomological Inoculation rate, ESTEIR/y= estimated entomological inoculation rate per year, ib/m/n= infective bite per man per night, b/p=number of mosquito bites per person, ib/m/y= infective bites per man/year

### Blood meal analysis in sampled malaria vectors

A total of 15 blood-fed resting adult *An. gambiae* s.l. caught during the dry (Allorkpem n=3, Pediatorkope n=2 and Tuanikope n=1) and rainy season (Allorkpem n=0, Pediatorkope n=1 and Tuanikope n=8) were analyzed to determine their blood meal sources. Out of these, 60% (9/15) fed on humans, 33.3% (5/15) fed on goats and pigs and 6.67% (1/15) neither fed on humans, goats or pigs. It was identified that 80% of the HBI was detected from indoor collected mosquitoes. The primary source of animal blood meals was identified as goats. A mixed blood meal source (human and goat) was identified from a single indoor mosquito. Additionally, mixed pig and goat bloodmeal were found in one indoor and one outdoor mosquito. Among the sites, a higher indoor HBI was recorded in Tuanikorpe (83.3%), followed by Allorkpem (33.3)] and then Pediatorkope (0), (Table 4)

**Table 4:** Blood meal sources of resting *An. gambiae* s.l. per site.

| Site | Blood-meal origins | *An. coluzzii* |  |
| --- | --- | --- | --- |
|  |  | Indoor | Outdoor |
| Allorkpem | Number Tested | 3 | 0 |
|  | Human | 1 (33.3) | 0 |
|  | Goat | 2 (66.7) | 0 |
|  | HBI | 33.3 | 0 |
|  | ABI | 66.7 | 0 |
| Pediatorkope | Number Tested | 2 | 1 |
|  | Human | 0 | 0 |
|  | Goat | 1 (50) | 1 (100) |
|  | Un-identified | 1 (50) | 0 |
|  | HBI | 0 | 0 |
|  | ABI | 100 | 100 |
| Tuanikope | Number Tested | 6 | 3 |
|  | Human | 3 (50) | 1 (33.3) |
|  | Goat | 1 (16.7) | 1 (33.3) |
|  | Human+Goat | 1 (16.7) | 0 |
|  | Human +Pig | 1 (16.7) | 1 (33.3) |
|  | HBI | 83.3 | 66.7 |
|  | ABI | 50 | 66.7 |

HBI: Human blood index, ABI: Animal blood index

### Insecticide resistant mutations in *Anopheles gambiae* s.l

The Vgsc-1014S and Vgsc-1014F mutations were genotyped in 845 *An. gambiae* s.l.. Another 300 *Anopheles gambiae* s.l. samples were genotyped for the presence of ACE-1 and Vgsc-1575Y mutations. Low frequencies of all mutations assayed were observed in the study sites (Table 5). However, the *kdr* mutation N1575Y had a relatively higher allelic frequency in Allorkpem (0.28) followed by Tuanikope (0.21) and then Pediatokope (0.07), (Table 5).

**Table 5:**
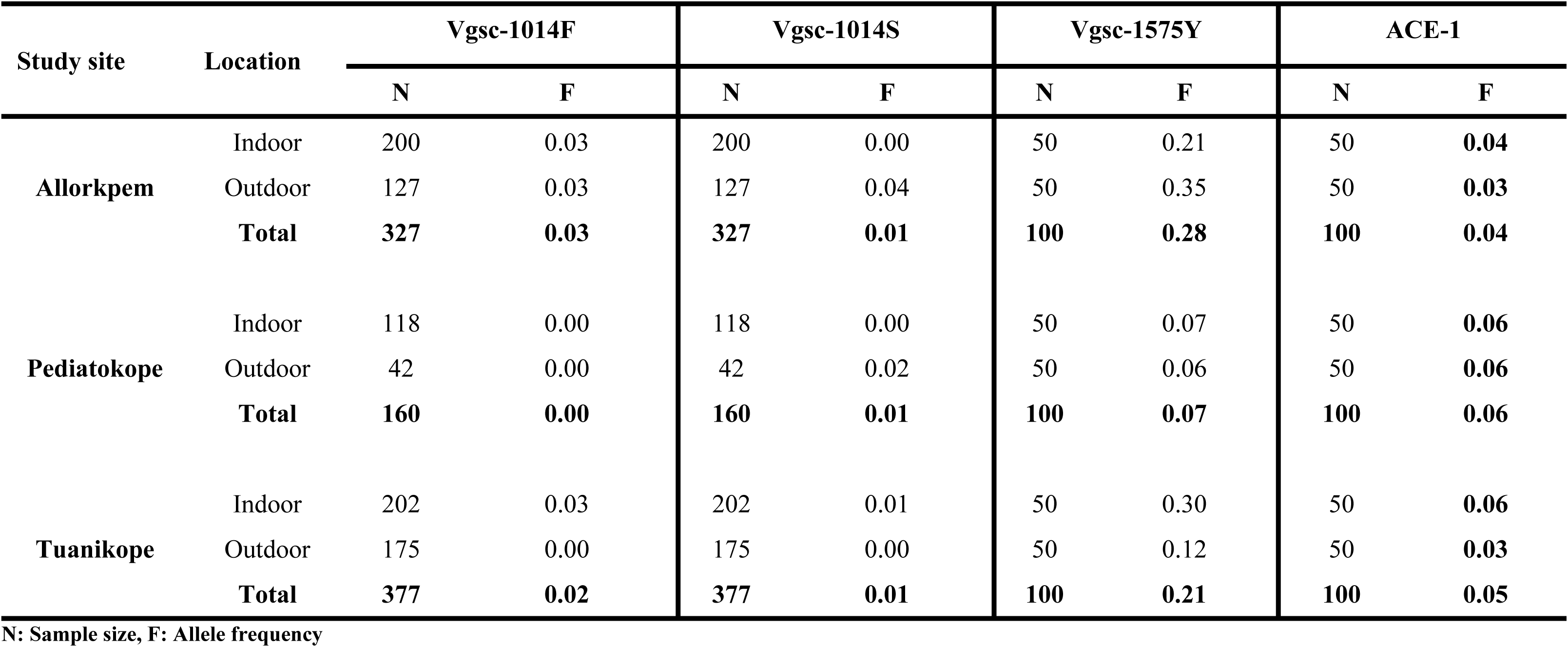
Frequency distribution of resistance genes in *Anopheles* mosquitoes collected from Island communities along the Volta Lake in Southern Ghana.

## Discussion

Understanding the bionomics, spatiotemporal distribution, and behavioral changes of malaria vectors in Island communities, is essential for designing targeted vector control measures. This study investigated the biting and feeding behavior, rates of infection with *Plasmodium falciparum* sporozoites, blood meal sources, and frequencies of insecticide resistance alleles of mosquitoes collected from Allorkpem, Pediatorkope, and Tuanikope. Higher densities of *Anopheles* mosquitoes were collected during the dry season. A high sporozoite rate with a low prevalence of *kdr* mutations was observed in *Anopheles* mosquitoes.

*An. gambiae* s.l. was the predominant vector for malaria transmission in Island communities year-round. However, its abundance exhibited significant seasonal variation. The abundance of *Anopheles gambiae* s.l. was significantly higher during the dry season compared to the rainy season. This is likely attributable to the windy climate during the rainy season at the sites. Contrary to this, several studies conducted in Ghana have reported that malaria vectors are abundant during the rainy season [3,20,27]. Previous studies by Monroe *et al* [28] in Zanzibar, an Island in Tanzania and Olanga *et al* [10] in Rusinga Island, Western Kenya also reported a high abundance of *Anopheles* mosquitoes in the rainy season.

Malaria vectors have been efficient in transmitting malaria mainly due to their anthropophilic and endophilic behaviour. Therefore, knowledge on the biting behavior of malaria vectors in local vector populations is important for deploying effective control tools and interrupting disease transmission. In this study, the abundance of *Anopheles* mosquitoes biting and resting indoors was higher than outdoors, in the dry and rainy seasons. Studies on the mainland in Ghana [20,29,30], however, observed an abundance of *Anopheles* mosquitoes biting outdoors. The high indoor biting and resting mosquitoes may be attributed to the absence of key vector control interventions, such as LLINs and IRS. Long-Lasting Insecticide Nets and IRS are cornerstone strategies in malaria prevention, effectively reducing mosquito densities and malaria transmission by targeting endophilic (indoor-resting) and anthropophilic (human-biting) vectors[31–33]. Their absence likely creates an unprotected environment that facilitates the abundance of mosquitoes within households. Moreover, outdoor biting mosquitoes could maintain residual malaria transmission [20]. Hence, continuous monitoring of mosquito behaviour is crucial to developing malaria control interventions.

This study revealed a concerning behaviour in *An. gambiae* s.l. mosquitoes: both indoor and outdoor populations preferred biting late at night, coinciding with human sleep times and maximizing their ability to spread malaria. The observed peak biting activity during the classical biting times also raises concerns, as this coincides with the dawn activities of the inhabitants, as residents wake up as early as 3:00 AM to fish, fetch water and firewood, and tend to animals. This creates a window of high vulnerability for malaria transmission and may potentially hinder control efforts in the region. [34].

An interesting observation from this study was the presence of other secondary malaria vectors: *An. rufipes* and *An. pharoensis*. These vectors have been implicated in other areas in transmitting malaria [35–37]. While their specific contribution to malaria transmission in Ghana remains to be fully elucidated, their presence emphasizes the complex nature of malaria transmission in the study area. Hence, the need for a comprehensive vector control strategy.

Sporozoite rates were observed to be relatively similar in both seasons across all study sites. Nonetheless, sporozoite-positive *Anopheles* mosquitoes collected indoors were more than those collected outdoors. This finding suggests sustained malaria transmission throughout the year, irrespective of seasonal changes. The higher proportion of sporozoite-positive *Anopheles* mosquitoes collected indoors compared to outdoors is particularly concerning, given the absence of indoor vector control tools such as LLINs and IRS. Without these interventions, indoor environments remain unprotected, allowing mosquitoes to thrive, feed, and rest. This may lead to a higher likelihood of transmission within households, implying a consistent level of malaria transmission throughout the year [14,38].

The entomological inoculation rate (EIR) remains the tool of choice for assessing malaria transmission intensity and endemicity [39,40]. Among the study sites, Tuanikope recorded the highest EIR, attributable to its high human biting rate (HBR) and sporozoite rate prevalence. Comparatively, other studies conducted in rural and urban areas across Africa have reported annual EIRs ranging from 0 to 884 and 0 to 43 infectious bites per person respectively [41,42]. The findings of this study highlight high malaria transmission in island communities, where residents lack adequate vector control measures and may experience relatively high annual EIRs (37.40 to 100.08) across all study sites. These values significantly exceed the EIR of 21.9 previously reported in Ghana’s coastal forest zone [43], although they remain below the EIR of 163 reported on Bioko Island, Guinea Bissau [18]. These data emphasize the urgent need for enhanced malaria control interventions in these vulnerable island populations.

Analysis of blood meals indicated a significant portion of malaria vectors displaying anthropophagic behavior. The increased proportions of infected *Anopheles gambiae* s.l. mosquitoes resting indoors raise concerns for malaria elimination initiatives, given the effectiveness of these mosquitoes in transmitting malaria [27,44,45].

The observed low frequency of resistance mutations in outdoor and indoor biting mosquito populations could be attributed to the absence of vector control tools in these areas, which could limit mosquito exposure to insecticides. During the study it was realized that no vector control campaigns has been done in the area for close to a decade, probably due to the hard-to-reach nature of the study sites. The rest of the country has LLIN distribution campaigns at least every two years. Without interventions such as IRS or LLINs, vectors may not be subjected to lethal or sub-lethal doses of insecticides, including those from alternative sources like aerosol sprays or agricultural pesticides [46]. This lack of exposure, combined with other factors influencing insecticide resistance, may contribute to the observed low allelic frequencies of resistant mutations. This observation suggests that the majority of the mosquito population in the study sites may be susceptible to insecticides used for vector control. Though the mutation frequencies were low, it raises concerns about controlling mosquito populations particularly those that spread diseases indoors, potentially worsening malaria outbreaks on the island.

## Conclusion

The findings of this study indicate that *Anopheles* species exhibited a seasonal abundance pattern, with higher densities during the dry season. A higher indoor biting and resting behavior was observed, although outdoor biting was evident, highlighting the need for complementary vector control strategies. Blood meal analysis revealed a strong anthropophilic feeding pattern for *Anopheles gambiae* s.l.. A high sporozoite rate with a low prevalence of resistant mutations was observed. These findings necessitate a comprehensive approach to vector control in the Island communities.

## Acknowledgment

Special appreciation to the entire inhabitants within our study sites, Christodea Haizel and Sebastian kow Egyin Mensah of the Centre for Vector-Borne Disease Research, Department of Medical Microbiology, University of Ghana Medical School.

## Availability of data and materials

All the data supporting this study are included in the article.

## Ethics approval and consent to participate

This study was approved by the Ghana Health Service ethics and Review Committee (GHS-ERC number: GHS-ERC: 051/03/23). Meetings were conducted at each study site with the chiefs, community leaders, and residents to introduce the research. Permission to conduct the study at the various sites was obtained from the community leaders. Prior to training and mosquito sampling, all adult volunteers participating in the HLC study provided written informed consent. Volunteers received a copy of their signed consent form, while another copy was securely stored in a locked cabinet within the Department of Medical Microbiology at the University of Ghana Medical School.” Verbal consent was obtained from household heads to sample mosquitoes in their houses and compounds.

## Consent for publication

Not applicable

